# High-resolution spatial transcriptomics of adult and pediatric human liver with Visium HD

**DOI:** 10.64898/2026.02.17.706350

**Authors:** Faizan Hasan, Rachel D. Edgar, Jawairia Atif, Diana Nakib, Cornelia Thoeni, Amanda Ricciuto, Blayne A. Sayed, Ian McGilvray, Gary D. Bader, Sonya A. MacParland

## Abstract

The liver is composed of diverse cell populations that coordinate essential metabolic and immune functions. Single-cell transcriptomics has advanced characterization of liver cellular composition, but dissociation of tissue to single-cells can introduce biases through the enrichment or depletion of cell types. Spatial transcriptomics is a complementary approach to avoid inherent bias for cell populations and to add important spatial context. The Visium HD spatial transcriptomics technology from 10X Genomics enables high-resolution spatial mapping of gene expression in tissue samples with a bin width of 2µm enabling quantification of transcripts at a sub-cellular resolution. We applied Visium HD to three healthy human liver donor samples, from two adult and one pediatric donor. We identified cell types by clustering 8µm bins and integration with single-cell reference maps. Differential expression analyses identified spatially distinct gene expression resulting in development of a high-resolution map of the liver. This resource provides cell-level and spatially-resolved insights into the cellular and anatomical heterogeneity of the liver to serve as a resource for researchers to identify disease-specific spatial signatures and novel therapeutic targets.

## Background and Summary

The liver plays a critical role in metabolism and immune function. These crucial roles are diminished in chronic liver diseases, leading to over two million deaths annually worldwide due to liver failure^1^. Single-cell transcriptomics has provided insights into the cellular composition of livers in health and disease, but is biased due to cell type-specific enrichment or depletion during the single-cell dissociation process. Previous work has highlighted difficulties in capturing specific populations such as cholangiocytes and hepatocytes^2^. In the cells captured, the process of dissociation has been shown to impact gene transcription^3^.

Spatial transcriptomics is a promising approach that does not have inherent bias for cell populations, and adds important spatial context. Spatial transcriptomics has been shown to capture important patterns of zonation in the human liver^4^, a dynamic feature important for understanding basic liver functions and alterations in liver disease processes. Until recently, spatial transcriptomic technologies have been low-resolution (Visium, 55µm) potentially mixing signals from multiple cells and multiple cell types. The latest spatial transcriptomic technology from 10X Genomics, Visium HD, enables high-resolution (2µm) spatial mapping of gene expression in tissue samples, offering a sophisticated platform for exploring the cellular composition of the liver.

Here we assayed samples from three neurologically deceased donor livers, with no evidence of histopathological liver disease. This data is at higher resolution than previous whole-transcriptome spatial datasets of healthy human liver^5^ enabling the identification of rare and transcriptionally distinct cell populations within the liver zonation structure. We provide these data as a resource for better understanding of the cellular and spatial liver heterogeneity.

## Methods

Figure 1A provides an overview of the liver spatial transcriptomics data generation which involves the collection of human liver tissue, preparing tissue sections, and Visium HD sample processing.

**Figure 1.**
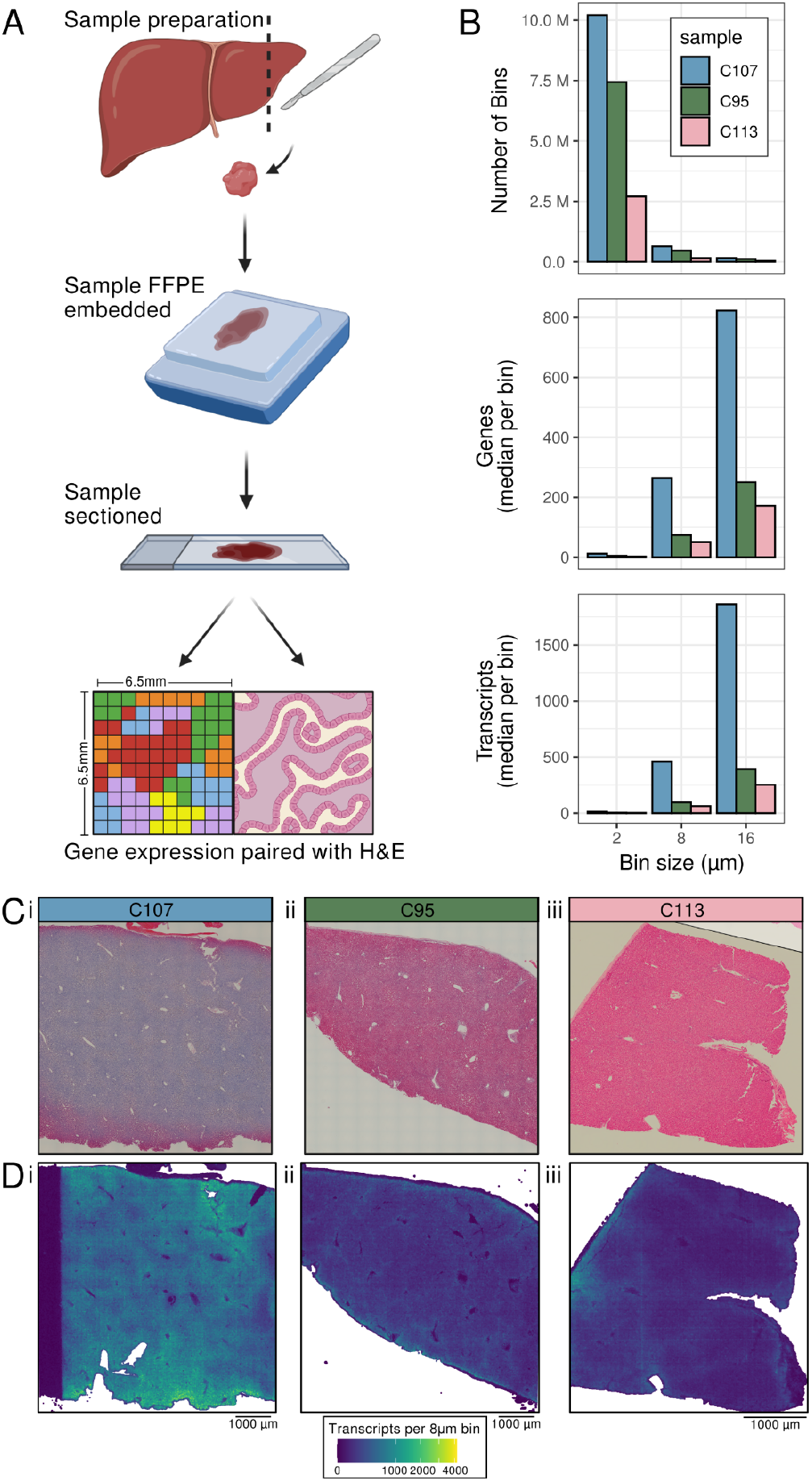
Sample processing of human liver for Visium HD resulted in three sample of varying quality. A) Sample processing workflow. B) Quality metrics of three samples at three bin sizes. C) H&E staining of samples processed with Visium HD. D) Location of 8µm bins coloured by number of UMI. Low quality areas in Di indicated with arrow.

### Human Liver Sample Collection

Healthy human liver tissue from the caudate or right liver lobe was obtained from neurologically deceased donor livers (**Table 1**). Donated livers had no evidence of histopathological liver disease supported by the hematoxylin and eosin (H&E) staining of the samples (**Supplementary data**).

**Table 1.** Spatial transcriptomic sample information.

| Sample | Age | Sex | Tissue | Storage | Reads | Sequencing saturation | Spacer Ranger version | Associated data previously published |
| --- | --- | --- | --- | --- | --- | --- | --- | --- |
| C107 | 51 | M | Caudate | Stored for 1 year at 4°C | 929,681,741 | 58.9% | 3.0.0 | None |
| C95 | 47 | F | Caudate | Stored 2 years; 1 year room temperature 1 year 4°C | 333,465,658 | 81.0% | 3.0.0 | Xenium spatial<sup>6</sup> |
| C113 | 15-19 | M | Right lobe | Stored for 2 years at 4°C | 225,446,253 | 93.1% | 3.0.1 | Dissociated single-cell<sup>6</sup> |

Fresh human liver samples were collected at the Hospital for Sick Children and Toronto General Hospital transplant centres. Liver tissue was prepared for Visium HD spatial transcriptomics by first processing into formalin-fixed, paraffin-embedded (FFPE) blocks which were then sectioned into 5µm thick slices and mounted onto 6.5mm by 6.5mm Visium HD slides (Visium HD v1 - FFPE; probe set v2.0) following the Visium HD Gene Expression User Guide.

Samples were sequenced on an Illumina NovaSeq X system. The Visium HD spatial transcriptomic data were sequenced to the depths between 300 - 950 million reads (**Table 1**). As sample C107 was initially only sequenced to a saturation of 34.4% the library was sequenced again for a final saturation of 58.9%. Reads were mapped to the human transcriptome (GRCh38-2020-A) and expression was quantified using Space Ranger software (**Table 1**). One sample (C113) was part of a tissue microarray (TMA) with three diseased liver samples included in the Visium HD capture area. Those diseased samples are not included in the sequencing data presented here, but are present in image files to enable future re-processing of the samples if needed.

### Ethics Statement

Healthy adult samples were collected with institutional ethics approval from the University Health Network (REB# 14-7425-AE) and the healthy pediatric sample through the Hospital for Sick Children (REB# 1000064039). All research was conducted in accordance with the Declarations of Helsinki and Istanbul, and written informed consent was obtained from living participants or, for pediatric or deceased donors, from their legal guardians or representatives.

### Annotation of Cell Type in 8µm Bins

Of the three samples processed, C107 had markedly higher quality, with twice as many genes and transcripts captured per bin (**Fig. 1B-D**). With only a few samples, it is uncertain why there was such variability in quality, however, cell annotation was performed on the highest quality sample C107.

To annotate bins with a cell type, an existing liver single-cell RNA-seq reference was mapped onto the spatial data using the Robust Cell Type Decomposition (RCTD)^7^ software for estimating cell type mixtures in spatial spots using a single-cell RNA-seq reference. To limit technical artefacts, low-quality bins with fewer than 100 UMIs were filtered out (this removed 22% of all bins for C107 which were mainly around the edges and in the lumen **Fig 1Di**) before cell type annotation of bins (**Fig 2C**). Spot class labels were assigned to bins using RCTD. The RCTD spot class “*singlet*” refers to bins predicted to contain only one cell type, whereas “*doublet-certain*” and “*doublet-uncertain*” refer to bins predicted to contain two cell types. Out of the high-quality bins, 67% were assigned an RCTD spot class and a cell type annotation (**Fig 2BC and Fig 3**). For the 33% of bins not well mapped to a cell type in the single-cell reference (i.e. RCTD “*rejects*”), these bins were clustered spatially using BANKSY^8^ spatially aware clustering followed by manual annotation of distinct cell types using known cell type markers and spatial context (**Fig. 3 and 4**).

**Figure 2.**
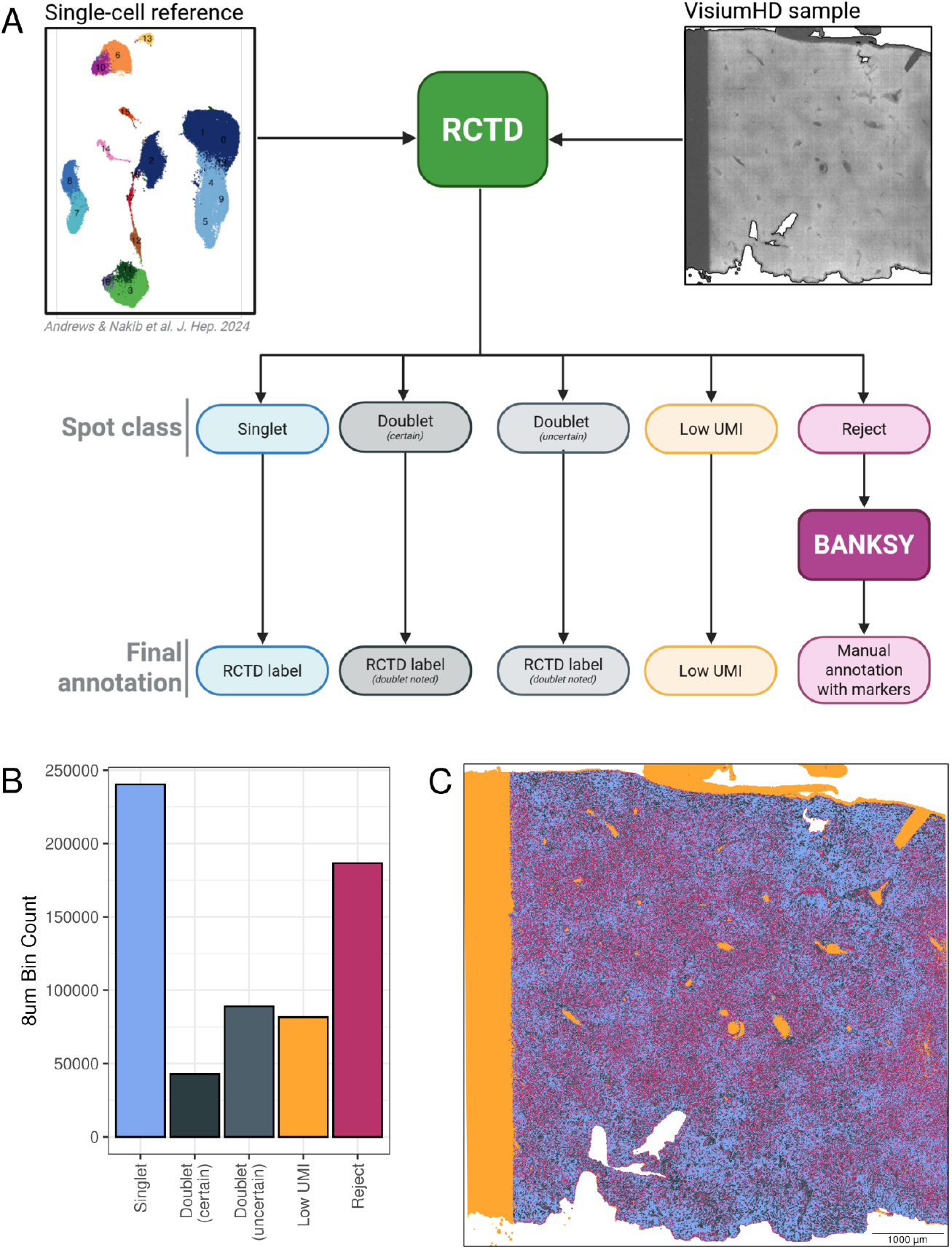
Annotation process for 8µm bins of sample C107. A) Workflow for cell annotation based on RCTD and BANKSY clustering when bins were rejected by RCTD. B) Spot class output of RCTD. C) Location of 8µm bins coloured by RCTD spot class.

**Figure 3.**
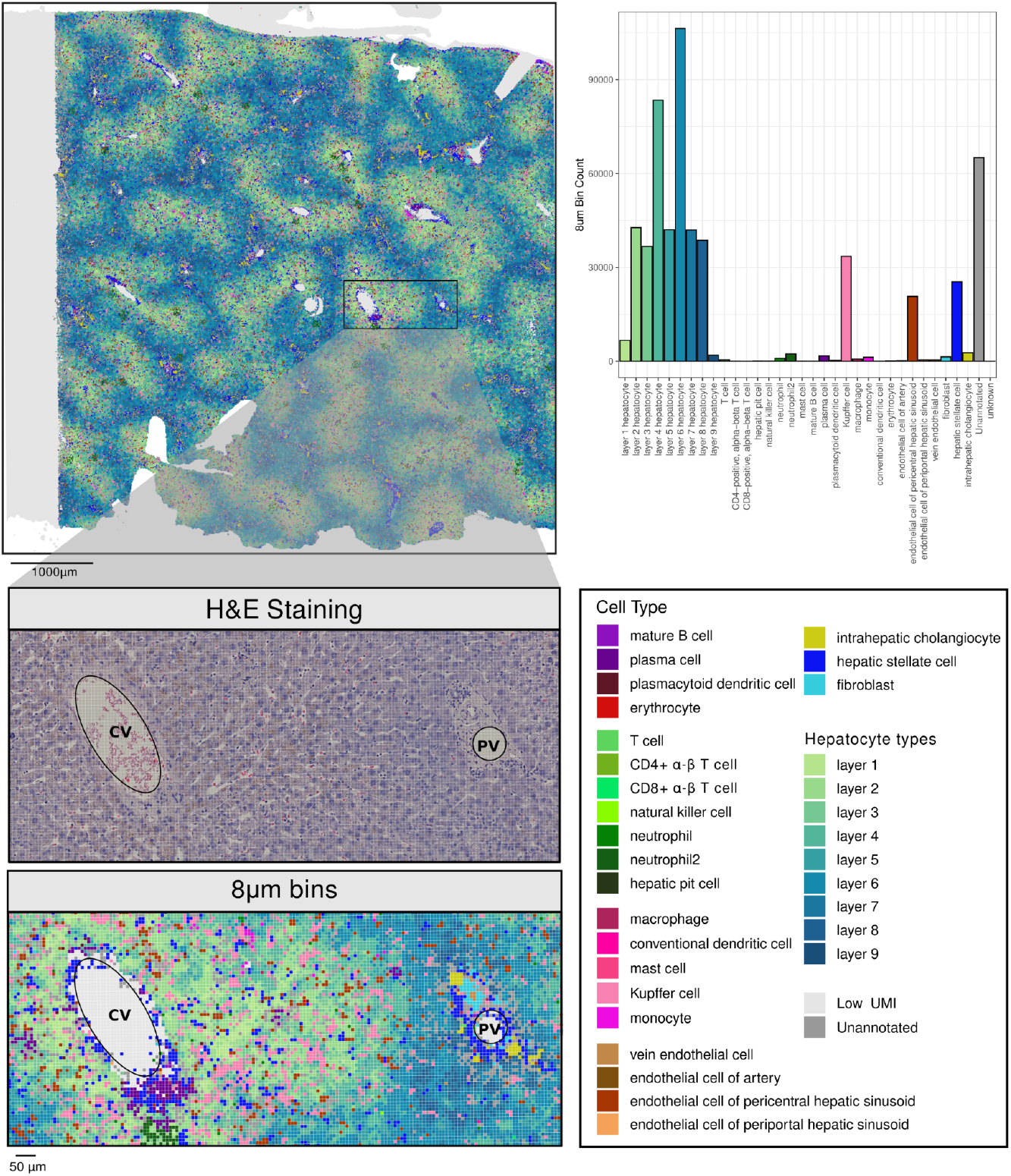
Expected cell types of the human liver can be captured, in their expected proportions and locations, with the Visium HD. A) Cell type annotation of the 8µm bins. B) Zoom in of a representative central vein (CV) to portal vein (PV) axis. C) Associated H&E staining of the same area. D) Counts of 8µm bins across the annotated cell types.

**Figure 4.**
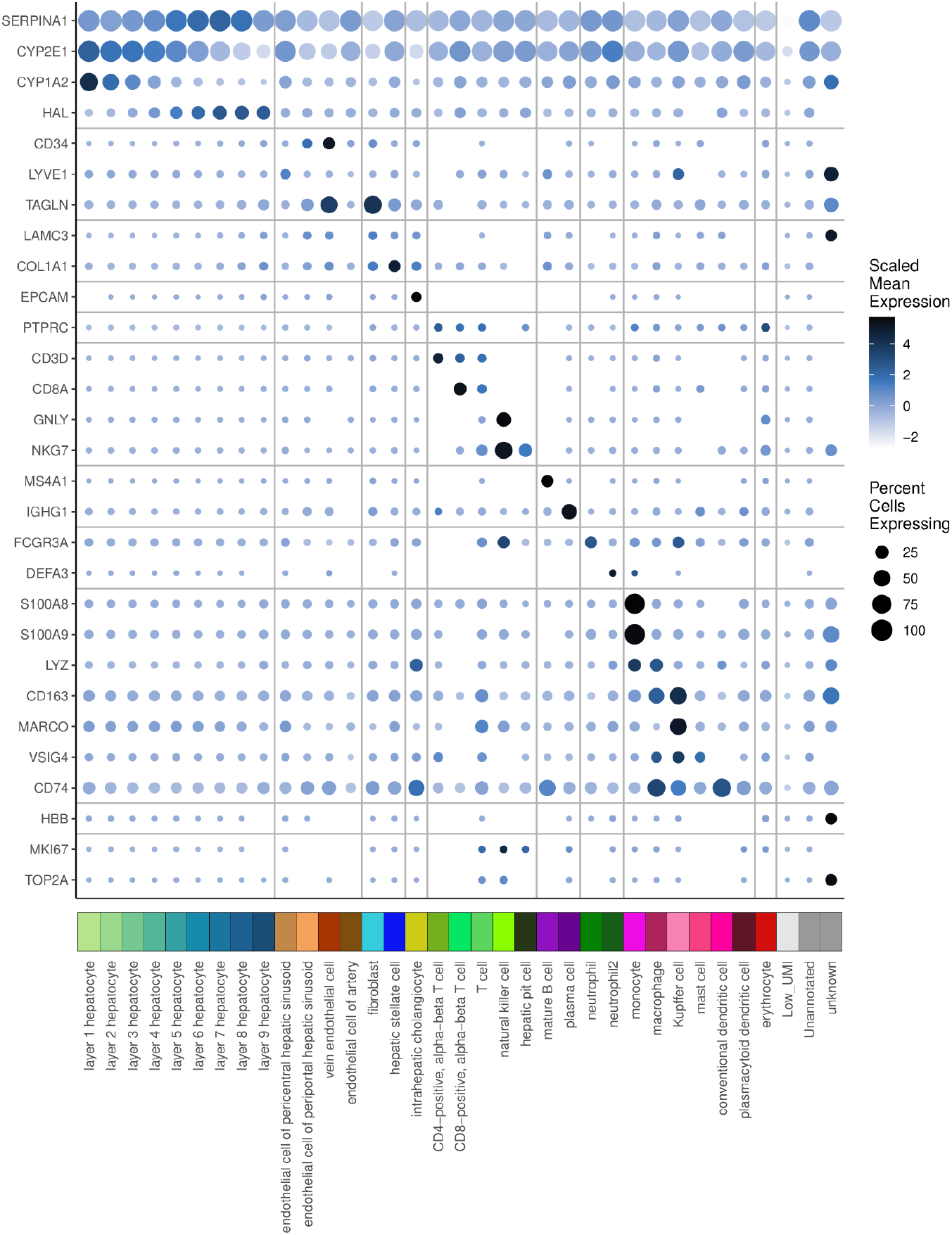
Expression of markers validates the cell types labels from cell annotation pipeline. Gene expression of key markers used for cell type annotation. The color of the points represents the gene expression level and size represents percent of cells of a type expressing the gene at all.

Liver tissue has defined spatial structure^9^ with hepatocytes arranged along a portal-to-central axis that generates metabolic and transcriptional gradients, a phenomenon known as liver zonation. We constructed a liver zonation axis in this data by dividing all bins into nine zonation layers, using a previously described method^10^. In brief, hepatocyte zonation scores were calculated using established marker genes^10^ and the *AddModuleScore* function from Seurat^11–13^. Bins were stratified into nine layers based on these scores. Based on these zonation scores individual bins were split into nine zonation layers. This zonation annotation was then smoothed by reassigning the zonation layer for a bin to the median zonation layer of its neighboring bins, as previously described^10^. Layer 1 corresponds to the most pericentral regions (near the central vein), layer 8 to the most periportal regions (near the portal triad), and layer 9 to regions more dense in fibroblasts and mainly but not exclusively periportal. For hepatocytes, the zonation layer was appended to the cell type name in the final annotation. Zonation of all genes across these layers was measured as differential expression between layers using the *FindAllMarkers* function from Seurat^11–13^ (with the default parameters except setting *logfc*.*threshold* = 0 and *min*.*pct*=0).

Pairing Visium HD transcriptional information and H&E images enabled the examination of structural features in the human liver. Lumen regions were identified in the H&E image based on colour intensity. The space in the image not covered by tissue (around the tissue section or in lumens) was identified by clustering the pixels of the H&E image by colour (k-means; k=5). The lightest coloured cluster captured this empty space. Then the empty space pixels were grouped spatially by distance from each other, using the graph_from_adjacency_matrix function in igraph^14^ to identify 110 lumen regions within the tissue. Lumen regions were classified by size to distinguish small sections of sinusoid (<150 pixels), small lumen (<600 pixels), and large lumen (>600 pixels). This translated to a mean diameter of 77µm for small lumen and 226µm for large. These lumen identified from the H&E image were overlaid with the 8µm bin coordinates to assign bins to lumen. Proximity to lumen regions was defined as: within 16µm (“adjacent”) and 80µm (“proximal”). Cell composition was then compared between regions adjacent or proximal to either small or large lumen. (**Fig. 5**).

**Figure 5.**
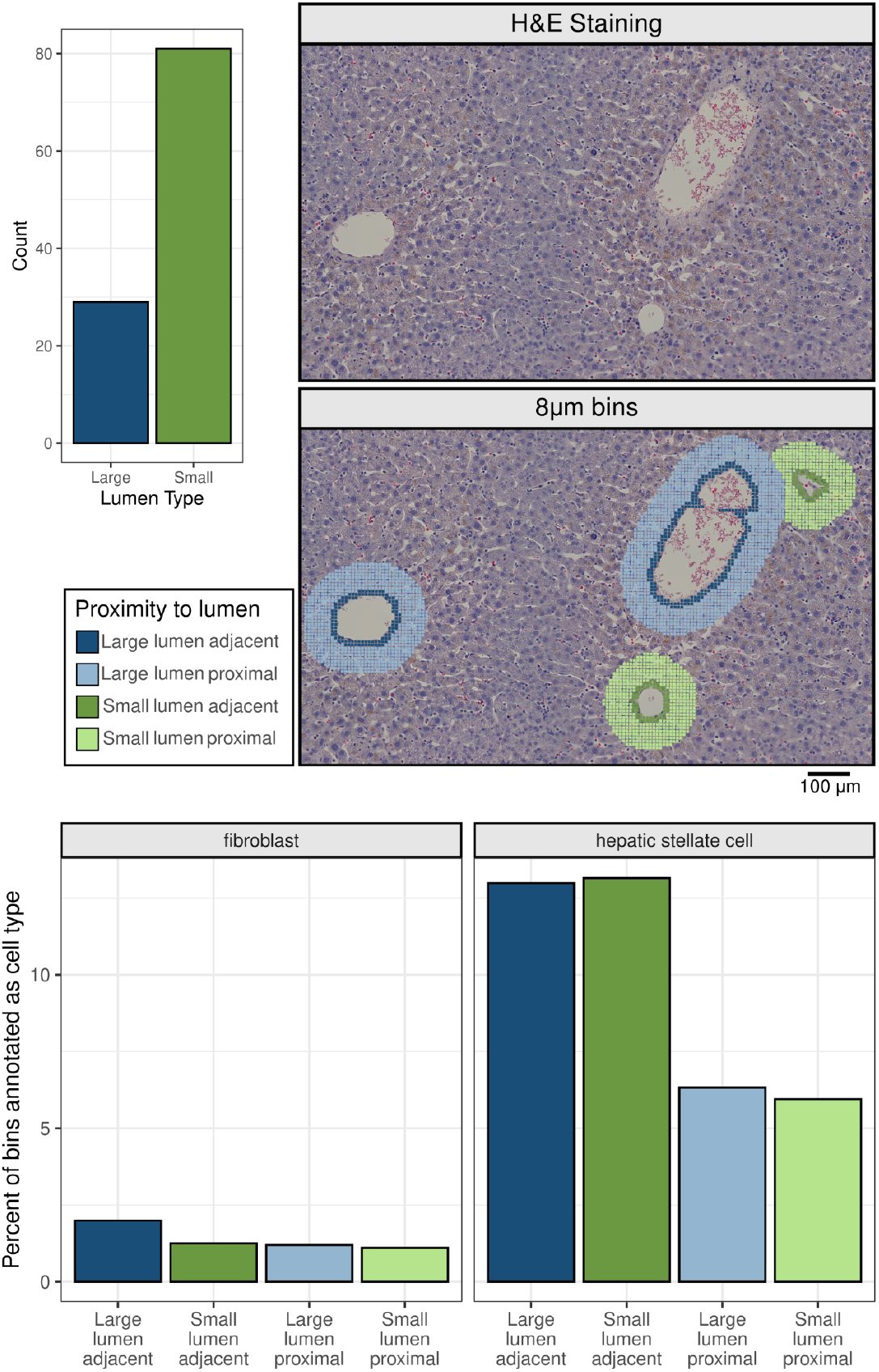
Lumen size can be quantified and associated with cell type composition. A) Distribution of lumen sizes among 110 lumen identified in the tissue. B) H&E staining of a representative region containing several lumens. C) Association of bins by proximity to lumen of two sizes at the 8um bin resolution. D) Percent of fibroblasts and hepatic stellate cells in regions adjacent (≤ 16 µm) or proximal (≤ 80 µm) to lumen across the entire tissue.

Cell annotation was performed at the 8µm bin resolution across the entire sample (**Fig. 1B-C**). To capture smaller cell types, at one representative central vein region, RCTD was run on 2µm bins with a cut off for low UMI of 20 (**Fig. 6**). At this central vein, cells lining the vein were annotated by RCTD as endothelial cells and these cells were visually distinct in the H&E images (**Fig. 6**). At this same central vein region, no 8µm bins at the lining of the veins were annotated as endothelial cells (**Fig. 6**). The ability to annotate the expected endothelial cells lining this vein demonstrates the utility of the 2µm bin resolution to identify smaller cells that may be missed at 8µm.

**Figure 6.**
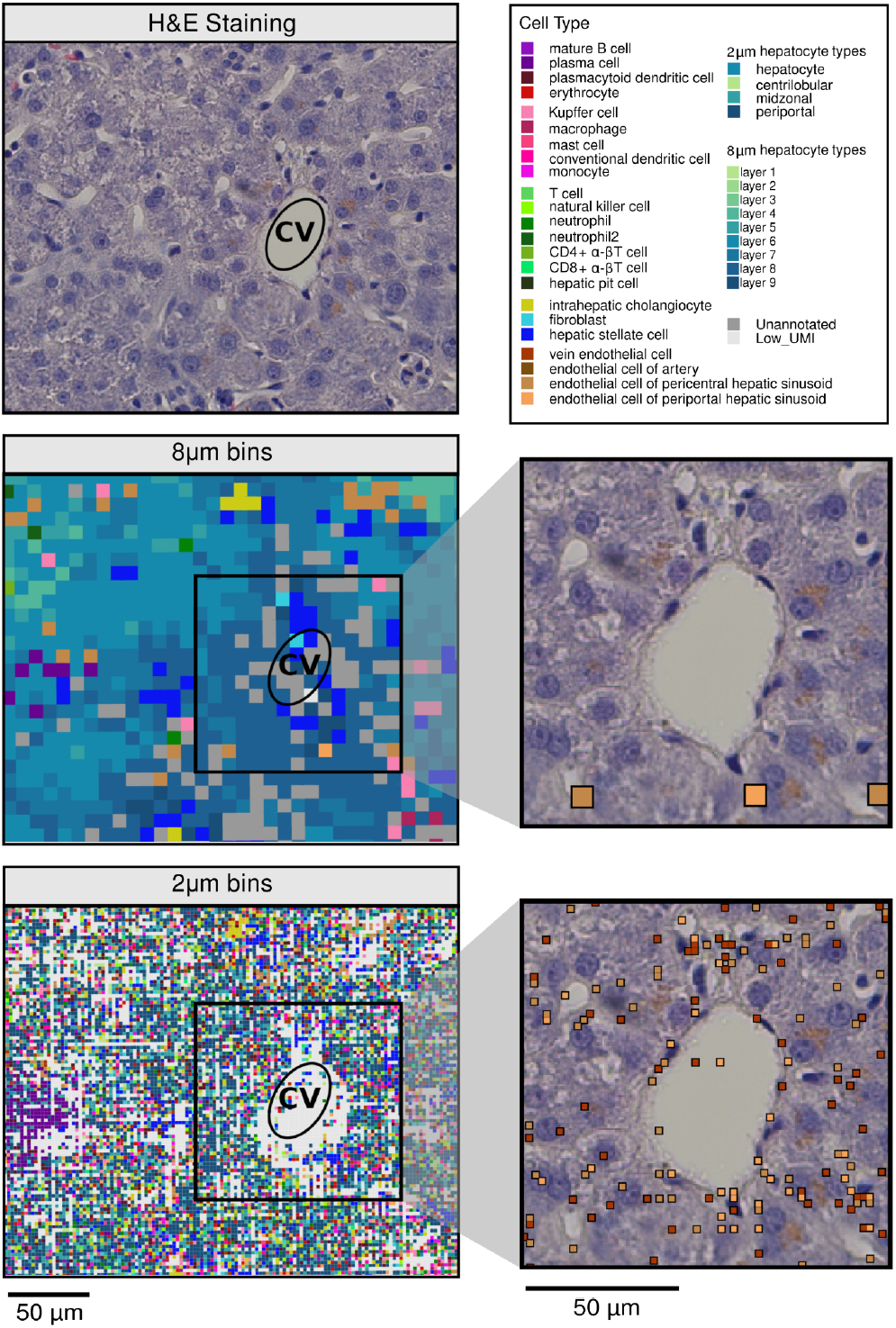
Though sparse in counts the 2µm bin resolution can capture endothelial lining at central veins not captured at the 8µm bin resolution. A) H&E staining of a representative central vein (CV) B) Cell type annotation at the 8um bin resolution. C) Further zoom in on the representative CV with only those 8µm bins annotated as endothelial highlighted. D) Cell type annotation at the 2µm bin resolution. E) Further zoom in on the representative CV with only those 2µm bins annotated as endothelial highlighted.Points are colored by endothelial cell type to highlight the distribution of vein endothelial cells (*TAGLN* and *SPARCL1* high) versus endothelial cell of pericentral hepatic sinusoid (*CLEC4G* and *LYVE1* high).

As bile ducts are made up of adjacent cholangiocytes, we identified bile ducts by grouping neighboring bins annotated as intrahepatic cholangiocytes. (**Fig. 7**). A cell–cell adjacency matrix, based on Euclidean distances between cholangiocytes, was converted into an undirected graph using the *graph_from_adjacency_matrix* function in igraph^14^. Connected groups of cholangiocytes were then identified within this graph using the *components* function, implementing a connected components algorithm, with default parameters. Of the 368 connected components of cholangiocytes, those containing more than 20 bins were classified as large bile ducts, those with 2–20 bins as small bile ducts, and groups with fewer than 2 bins were excluded as potential technical artifacts. This translated to a mean diameter of 26µm for small ducts and 82µm for large.

**Figure 7.**
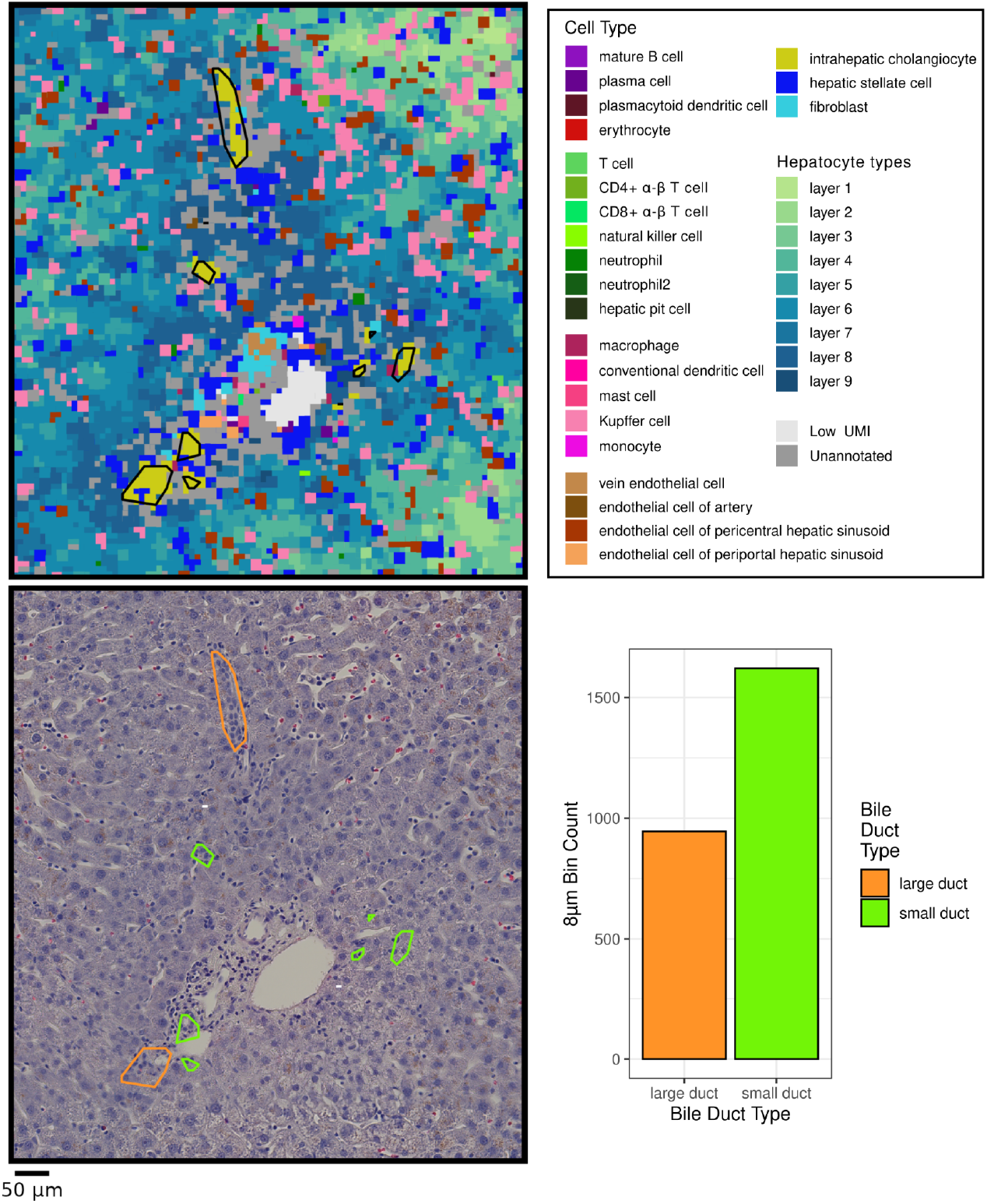
Bile ducts of different sizes can be identified. A) Cell type annotation at the 8um bin resolution at representative periportal region. Transcriptionally identified bile ducts are highlighted with a black outlines. B) H&E staining of representative region with transcriptionally identified bile ducts are highlighted and coloured by size type. C) Counts of cholangiocyte bins which were assigned to either small or large bile ducts.

### Validation of Gene Expression Measurements by Immunohistochemistry

Immunohistochemical images were obtained from the Human Protein Atlas (https://www.proteinatlas.org) and the annotation of central and portal veins were confirmed by a liver pathologist (CT). The representative immunohistochemistry (IHC) stain shown is for: SLCO1B3 - antibody HPA004943 patient id 1846; LGALS4 - antibody HPA031186 Patient id 1720.

### Data Record

The gene expression data, associated H&E images, annotation of the 8 µm bins and an interactive Shiny app for exploring regions of interest will be made publicly available upon acceptance of the manuscript. For two samples additional spatial and dissociated single-cell data have been published^6^.

### Technical Validation

To technically validate the data, we assessed its ability to capture a strong, expected biological signal. In liver spatial transcriptomics, one of the most prominent such signals is hepatocyte zonation. We therefore evaluated the ability of Visium HD toTo technically validate the data, we assessed its ability to capture a strong expected biological signal. In liver spatial transcriptomics, one of the strongest expected signals is hepatocyte zonation. We therefore wanted to validate the ability of the Visium HD to capture hepatocyte zonation. capture this pattern. Differential expression analysis of Visium HD bins revealed clear periportal and pericentral markers (**Fig. 8A**). Fold-change values were calculated by comparing all 8 µm bins assigned to a given zonation region against all other bins (**Fig. 8A**). The spatial distribution of these genes across the tissue section recapitulated the expected zonation patterns (**Fig. 8B**). To validate the ability of Visium HD to capture zonation, we examined both a well-established zonation marker (*SLCO1B3*) and a less well-characterized zonated gene (*LGALS4*). Importantly, protein-level validation using immunohistochemistry (IHC) data from the Human Protein Atlas confirmed the zonated expression of these markers (**Fig. 7C**), supporting the ability of Visium HD to capture physiologically relevant spatial organization in the liver.

**Figure 8.**
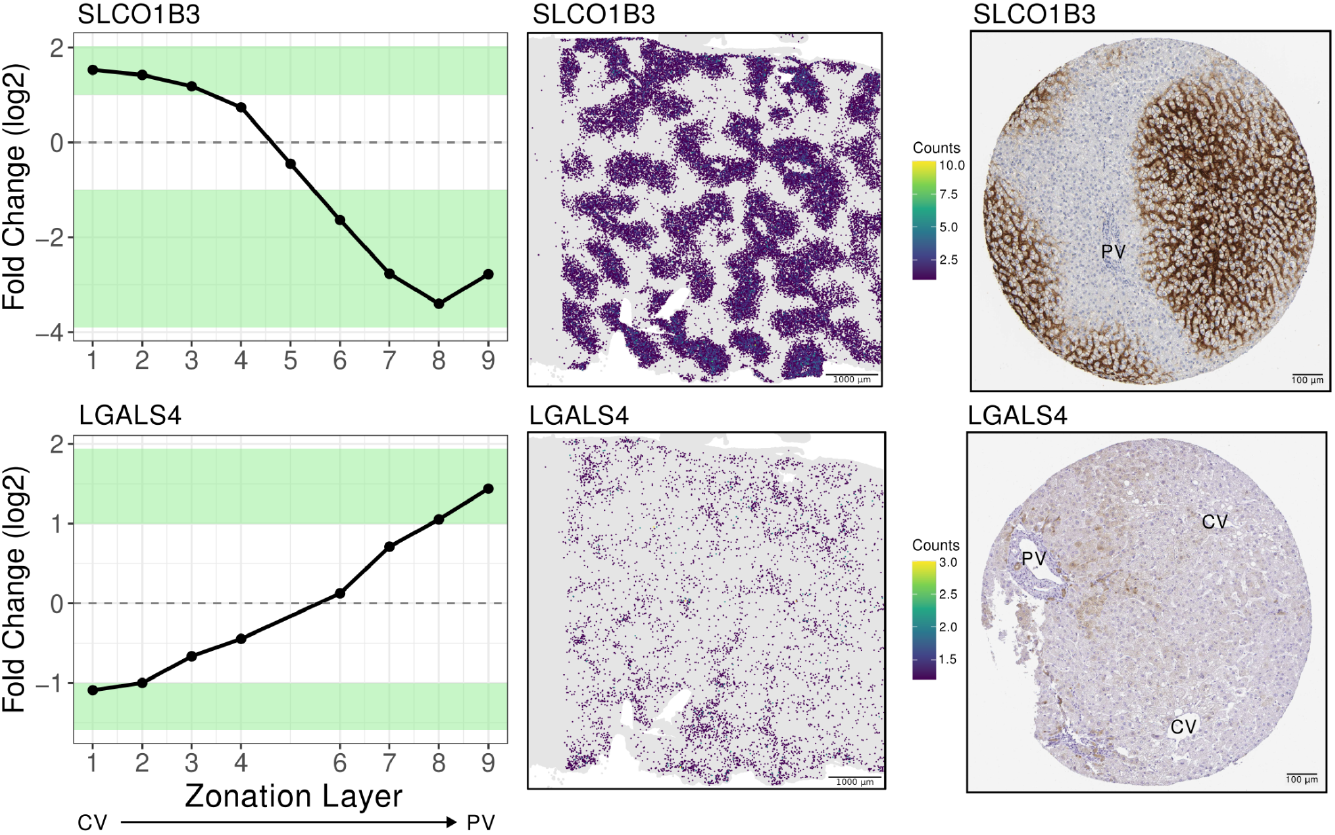
Markers of zonation can be identified in the Visium HD expression data and validated at the protein level. A) Differential expression fold change of representative zonated genes. Fold change values are comparing all 8µm Visium HD bins in a given zonation bin to all others. B) Expression across the entire sample. C) Representative IHC plots verifying the zonation of genes at the protein level. Annotations are central vein (CV) and portal vein (PV)

## Author Contributions

F.H., R.D.E., G.D.B., and S.A.M. conceived the study. F.H and R.D.E. performed data analysis. J.A., D.N., and C.T. contributed to investigation and data curation. A.R., B.S., I.M., G.D.B., and S.A.M. provided resources and supervision. F.H and R.D.E. drafted the manuscript with input from all authors. All authors reviewed and approved the final manuscript.

## Acknowledgements

The authors acknowledge the University Health Network Pathology Research Program and Princess Margaret Genomics Centre for their support and services. The concept figures were created with Biorender.com.

